# Diversity and seasonality of horse flies (Diptera: Tabanidae) in Uruguay

**DOI:** 10.1101/794479

**Authors:** Martín Lucas, Tiago K. Krolow, Franklin Riet-Correa, Antonio Thadeu M. Barros, Rodrigo F. Krüger, Anderson Saravia, Cecilia Miraballes

## Abstract

Horse flies (Diptera: Tabanidae) are hematophagous insects that cause direct and indirect losses in livestock production and are important vectors of pathogens. The aim of this study was to determine the diversity and seasonality of horse fly species at an experimental farm in Tacuarembó and the diversity of species in different departments of Uruguay. For 20 months, systematic collections were performed using Nzi and Malaise traps in two different environments at the experimental farm. Temperature, humidity and rainfall were recorded using a local climatological station. In addition, nonsystematic collections were made at farms located in the departments of Paysandú, Tacuarembó and Colonia. A total of 3,666 horse flies were collected, allowing the identification of 16 species. Three species were recorded for the first time in Uruguay: *Dasybasis ornatissima* (Brèthes), *Dasybasis missionum* (Macquart), and *Tabanus aff. platensis* Brèthes. A species that had not been previously taxonomically described was identified (*Tabanus* sp.1). In the systematic captures, the most abundant species were *Tabanus campestris* Brèthes, *T. aff. platensis* and *D. missionum*, representing 77.6% of the collected specimens. The environment was an important factor related to the abundance of horse flies, as well as the mean temperature. The horse fly season in Tacuarembó started in September and ended in May, with three evident peaks, the most important one during summer. No horse flies were caught during winter. Variations in the prevalence of species in the different departments were observed, indicating the need to carry out new sampling efforts in different areas.

## Introduction

Horse flies (Tabanidae) are hematophagous dipterans that cause direct losses to livestock production due to irritation, stress, and blood loss in animals, mainly in cattle and horses (Baldacchino et al. 2014). A decrease in weight gains of 0.1 kg up to one kilogram per day in cattle has been described due to the direct effect of horse flies (Foil and Hogsette 1994; Baldacchino et al. 2014). Economic losses are directly related to the number of horse flies present in the environment (Perich et al. 1986). In addition to the losses caused by the direct effects of horse flies, they cause indirect losses due to their role as a mechanical vector of numerous pathogens, including those causing bovine leukosis, vesicular stomatitis, equine infectious anemia, swine fever, anthrax, tularemia, various species of trypanosomes and *Anaplasma marginale* (Krinsky 1976; Foil 1989). These indirect losses may be even more important than the direct losses, but not all members of the Tabanidae family have the same potential for transmitting disease agents because their hematophagous behavior and anatomical characteristics, which determine the amount of blood they can transport, varies (Magnarelli and Anderson 1980; Foil 1989; Scoles et al. 2008).

Uruguay has a subtropical climate according to the Köppen classification, presenting four marked seasons with a mean annual temperature of 17.29 °C and a mean humidity of 76.03% (INIA 2019). It is included in the Pampa biome, along with the state of Rio Grande do Sul (Brazil) and the province of Buenos Aires (Argentina).

The emergence of the first generation of horse flies during the year depends on the latitude and the season (Chvála et al. 1972; Kruger and Krolow 2015). Horse flies are active mostly in warm seasons, when both the relative humidity and temperature are high. Only females are hematophagous, and adult longevity varies from two to three weeks (Baldacchino et al. 2014). Although the females need a blood meal every three or four days, their hematophagous habits are painful, and animals try to remove them, which may cause the interruption of feeding. This interruption causes females to seek other animals to complete a meal (Hornok et al. 2008); they are able to fly several kilometers, reaching speeds of five m/s, which highlights their potential as a mechanical vector (Hornok et al. 2008).

To study the diversity and abundance of horse fly species in different environments, a wide variety of traps have been used (Thorsteinson et al. 1965; Thompson 1969). However, the Nzi trap was designed specifically to catch biting flies (Mihok 2002), such as horse flies and stable flies (Stomoxyinae). This trap has been shown to capture a greater number of horse flies and species than other traps (Van Hennekeler et al. 2008), but some genera of horse flies tend to be captured in higher numbers by other traps (Baldacchino et al. 2014).

Approximately 4300 species belonging to 137 genera of horse flies have been described worldwide (Coscarón and Papavero 2009; Coscarón and Martínez 2019). The Tabanidae family is subdivided into four subfamilies: Chrysopsinae, Pangoniinae, Scepsidinae and Tabaninae. The subfamilies Chrysopsinae and Tabaninae are considered the most relevant in the mechanical transmission of pathogens (Hawkins et al. 1982; Scoles et al. 2008). In the Neotropical region, 71 genera and 1205 species have been described (Coscarón and Papavero 2009; Henriques et al. 2012; Coscarón and Martínez 2019). Regionally, 350 species of Tabanidae have been described in Argentina (Coscarón 1998) and 480 in Brazil (Taxonomic Catalog of Fauna of Brazil 2019).

In Uruguay, 43 species belonging to 14 genera have been reported (Coscarón and Martínez 2019); however, most of these captures were performed in the XIX and XX centuries, so it is unknown whether this diversity of horse flies still exists in the country. Additionally, there have been no studies in the country regarding the abundance, ecology and seasonality of the Tabanidae family.

The objectives of this study were to evaluate the seasonality of the horse fly species present at an experimental farm in Tacuarembó and the diversity of species in different departments of Uruguay.

## Materials and Methods

For 20 months, between October 2017 and June 2019, systematic collections were performed using four Nzi traps and two Malaise traps at the Experimental Farm of INIA La Magnolia (EFILM) in the department of Tacuarembó (31°42’29.0” S, 55° 48’09.1” W). Two Nzi traps and one Malaise trap were placed in a “lowland” habitat with native forest and creeks, while the other two Nzi traps and one Malaise trap were placed in “highland” habitat, which were clearer environments, near a cultivated eucalyptus forest. All these traps remained active during the 20 months of the study. According to a conventional climatological station located at the EFILM, the following experimental parameters were recorded on a monthly basis: average maximum, average minimum and average mean temperatures; average mean relative humidity and accumulative rainfall.

In addition, nonsystematic collections were performed on farms located at the departments of Paysandú, Tacuarembó and Colonia using traps and/or manually. These captures were carried out with the aim of expanding the diversity of horse fly species collected. A defined collection time pattern was not established, and the capture methods varied, so these data were not considered for the analysis of seasonality and abundance of the horse fly species.

Of the total number of horse flies captured, samples from one to 10 specimens of each suspected species were prepared for taxonomic identification.

## Data collection

### Systematic collections

Between October and December 2017, two Nzi traps (Rincon-Vitova, USA) (Nzi 1 and Nzi 4) and one Malaise trap (Rincon-Vitova, USA) (Mal 6) were placed in a clear field environment close to artificial eucalyptus forests (highland), and two Nzi traps (Nzi 2 and Nzi 3) and one Malaise trap (Mal 5) were placed in an environment close to creeks and native forest (lowland). Because alcohol was not used in the collectors for the conservation of horse flies, during summer (December to February), when the populations of horse flies were highest, the collections were carried out at least three times a week to prevent desiccation of the captured specimens. During winter (June to August), collections were performed at least once a week, while in spring (September to November) and autumn (March to May), the collections were made from one to three times a week depending on the number of specimens collected.

### Nonsystematic collections

To identify species in different areas of the country or species that might not be captured by traps, sporadic samples were performed in the departments of Tacuarembó (three farms), Paysandú (one farm) and Colonia (two farms) (Table 1). These samplings were performed manually and/or with Nzi traps. In the manual sampling, some horse flies were caught while feeding on animals, while others were manually caught in the environment.

**Table 1.**
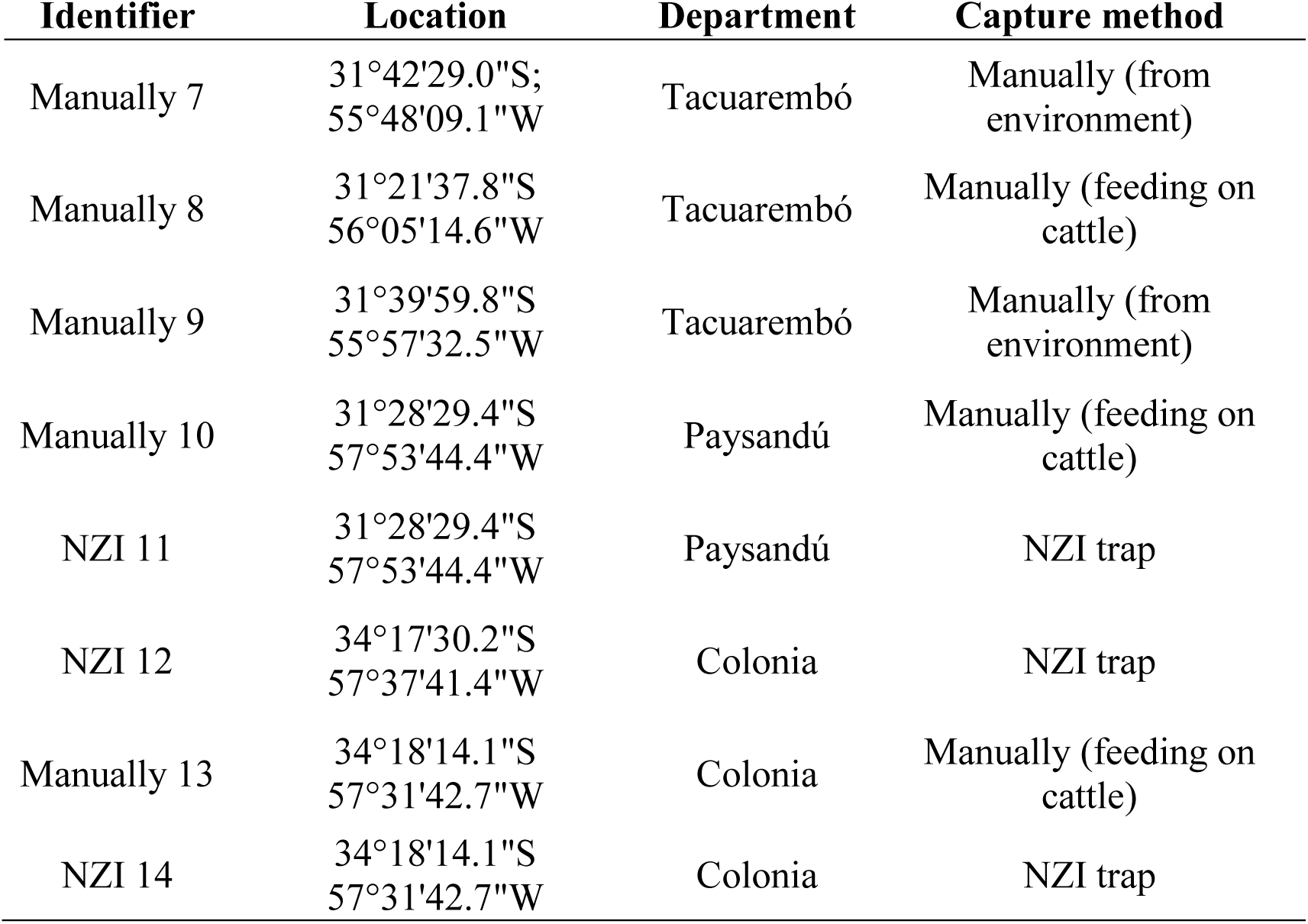
Location and capture method of nonsystematic horse flies collections.

| Identifier | Location | Department | Capture method |
| --- | --- | --- | --- |
| Manually 7 | 31°42'29.0"S;<br>55°48'09.1"W | Tacuarembó | Manually (from environment) |
| Manually 8 | 31°21'37.8"S<br>56°05'14.6"W | Tacuarembó | Manually (feeding on cattle) |
| Manually 9 | 31°39'59.8"S<br>55°57'32.5"W | Tacuarembó | Manually (from environment) |
| Manually 10 | 31°28'29.4"S<br>57°53'44.4"W | Paysandú | Manually (feeding on cattle) |
| NZI 11 | 31°28'29.4"S<br>57°53'44.4"W | Paysandú | NZI trap |
| NZI 12 | 34°17'30.2"S<br>57°37'41.4"W | Colonia | NZI trap |
| Manually 13 | 34°18'14.1"S<br>57°31'42.7"W | Colonia | Manually (feeding on cattle) |
| NZI 14 | 34°18'14.1"S<br>57°31'42.7"W | Colonia | NZI trap |

### Statistical analysis

To perform descriptive and inferential analyses, two databases were created in Excel, one for the systematic captures and the other for the nonsystematic captures and were then imported into STATA 2014 (Stata Corp 2015) after identifying any data errors.

The datasets generated during and/or analyzed during the current study are available at Miraballes et al (2019).

### Descriptive analysis

Descriptive analyses were performed based on the total number of individuals captured by location, date, season, trap type, type of manual capture (environmental or on animals) and species.

### Inferential analysis

The influence of mean temperature (mt) and environment (lowland and highland) on the abundance of horsefly species was tested using generalized linear models (GLMs). The models were obtained by extraction of significant terms (p<0.05) from the full model, which included all of the variables mt, maximum temperature, minimum temperature, relative humidity, rainfall, and environment and their interactions, as suggested by Crawley (2007). Each term was analyzed followed by ANOVA and chi-square tests (Chi) to recalculate the deviation explained by the other terms.

We tested the hypothesis that the mt in different environments alters the abundance of Tabanidae using models with response variables that assume integer values with regard to abundance. The explanatory variables that were used were mt and environment without interaction terms. The complete models that were used for hypothesis testing are as follows:

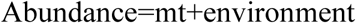

The different years in which the experiments were conducted were not used as explanatory variables because there was autocorrelation between mt and year. In the models, a plus sign (+) denotes the addition of a variable. A quasi-Poisson error distribution and log link function were used to model the estimated abundance.

## Results

During the study, 3,666 horse flies were collected: 3,211 through systematic collections and 455 by nonsystematic ones. These specimens represented two subfamilies, three tribes, six genera and 16 species. Fourteen species were definitively identified taxonomically, and one species was preliminarily identified as *Tabanus affinity platensis* Brèthes (*T. aff. platensis*); an undescribed species of *Tabanus* (*Tabanus* sp.1) was found. In addition, it was not possible to taxonomically identify 11 individuals belonging to the genus *Tabanus* that were likely members of the same species, and they were classified as *Tabanus* spp.

### Systematic collections

Of the 3,211 specimens captured by this method, a total of 15 species were identified, of which three were *Tabanus campestris* Brèthes; *T. aff. platensis* and *Dasybasis missionum* (Macquart) represented 77.6% of the total captures (Table 2). Other species of varying relative abundance identified were a new species to be described (*Tabanus* sp.1), *Tabanus triangulum* Wiedemann, *Tabanus acer* Brèthes and *Poeciloderas quadripunctatus* (Fabricius). The remaining species only accounted for 3.0% of the captures. Additionally, 3.4% of the individuals could not be identified because they were in poor condition.

**Table 2.** Number of individuals and species of horse flies caught by traps in systematic collections between October 2017 and June 2019

| Species | Lowland |  |  | Highland |  |  | TOTAL (%) |
| --- | --- | --- | --- | --- | --- | --- | --- |
|  | Traps |  |  |  |  |  |  |
|  | NZI 2 | NZI 3 | MAL 5 | NZI 1 | NZI 4 | MAL 6 |  |
|  | Number of individuals |  |  |  |  |  |  |
| *Catachlorops circumfusus* | 0 | 1 | 0 | 0 | 2 | 1 | 4 (0.1) |
| *Chrysops brevifascia* | 1 | 7 | 2 | 1 | 0 | 0 | 11 (0.3) |
| *Dasybasis missionum* | 96 | 564 | 21 | 2 | 14 | 18 | 715 (22.3) |
| *D. ornatissima* | 0 | 1 | 0 | 0 | 0 | 0 | 1 (0.0) |
| *Poeciloderas lindneri* | 0 | 17 | 0 | 0 | 0 | 1 | 18 (0.6) |
| *P. quadripunctatus* | 3 | 30 | 0 | 0 | 0 | 0 | 33 (1.0) |
| *Tabanus acer* | 9 | 31 | 1 | 6 | 12 | 3 | 62 (1.9) |
| *T. aff. platensis* | 3 | 837 | 0 | 14 | 11 | 2 | 867 (27.0) |
| *T. campestris* | 28 | 863 | 2 | 5 | 6 | 7 | 911 (28.4) |
| *T. claripennis* | 1 | 17 | 0 | 0 | 5 | 0 | 23 (0.7) |
| *T. fuscofasciatus* | 3 | 16 | 0 | 0 | 1 | 0 | 20 (0.6) |
| *T. fuscus* | 1 | 17 | 1 | 0 | 0 | 0 | 19 (0.6) |
| *T. pungens* | 0 | 0 | 0 | 0 | 0 | 1 | 1 (0.0) |
| *T. sp.1* | 59 | 203 | 3 | 23 | 40 | 14 | 342 (10.7) |
| *T. triangulum* | 5 | 24 | 3 | 27 | 14 | 3 | 76 (2.4) |
| Unidentified | 6 | 87 | 9 | 1 | 3 | 2 | 108 (3.4) |
| TOTAL (%) | 215 (6.7) | 2715 (84.5) | 42 (1.3) | 79 (2.5) | 108 (3.4) | 52 (1.6) | 3211 (100) |

Traps located in the “lowland” habitat were responsible for 92.5% of the total catches during the study, and within this habitat the Nzi 3 trap captured 84.5% of the total specimens (Table 2). The most abundant species in the “lowland” were *T. campestris* (30.0%), *T. aff. platensis* (28.3%) and *D. missionum* (22.9%). Captures in the “highland” habitat represented only 7.5% of the total and the three most abundant species were: *Tabanus* sp.1 (32.2%), *T. triangulum* (18.4%) and *D. missionum* (14.2%).

Figure 1 depicts the seasonality of the horse flies in the “lowland” habitat, where most individuals were caught, and shows that they were active from the end of September to the beginning of May, with no activity from June to August. It also shows the presence of three peaks: one at the end of spring, another one in mid-summer (the most pronounced one), and a third one during the fall. A decrease in the total captures during the second season was also observed.

**Fig. 1.**
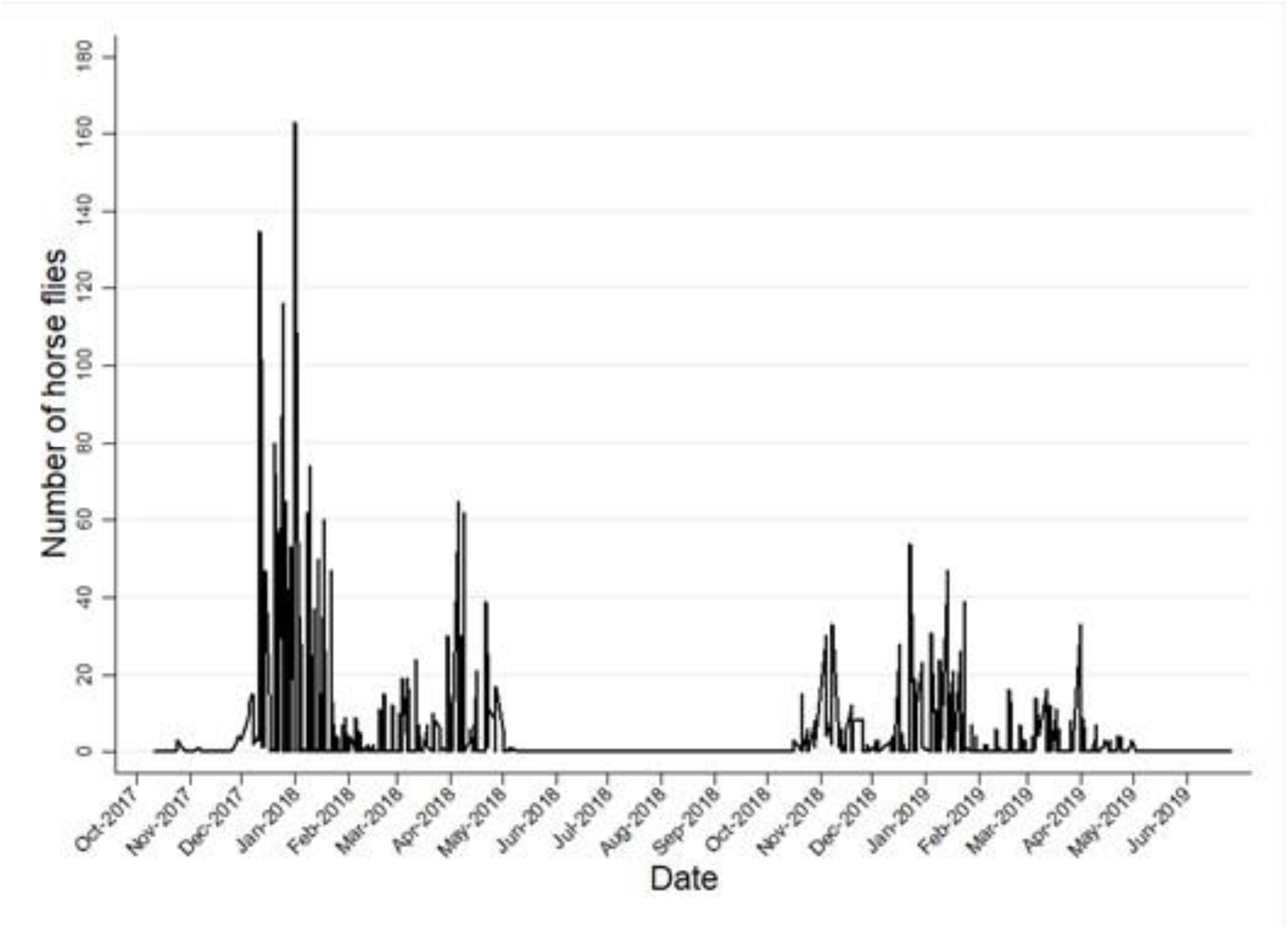
Number of horse flies caught in the “lowland” environment by systematic captures during the study period.

Of the total captures in both habitats, 2,336 individuals were captured during the first season of horse fly activity (September 2017-May 2018), while 875 horse flies were captured during the second season (September 2018-May 2019). Figure 2 shows the different seasonality of the three most prevalent species. It can be observed that *D. missionum* showed a different behavior, presenting a peak during late spring and autumn.

**Fig. 2.**
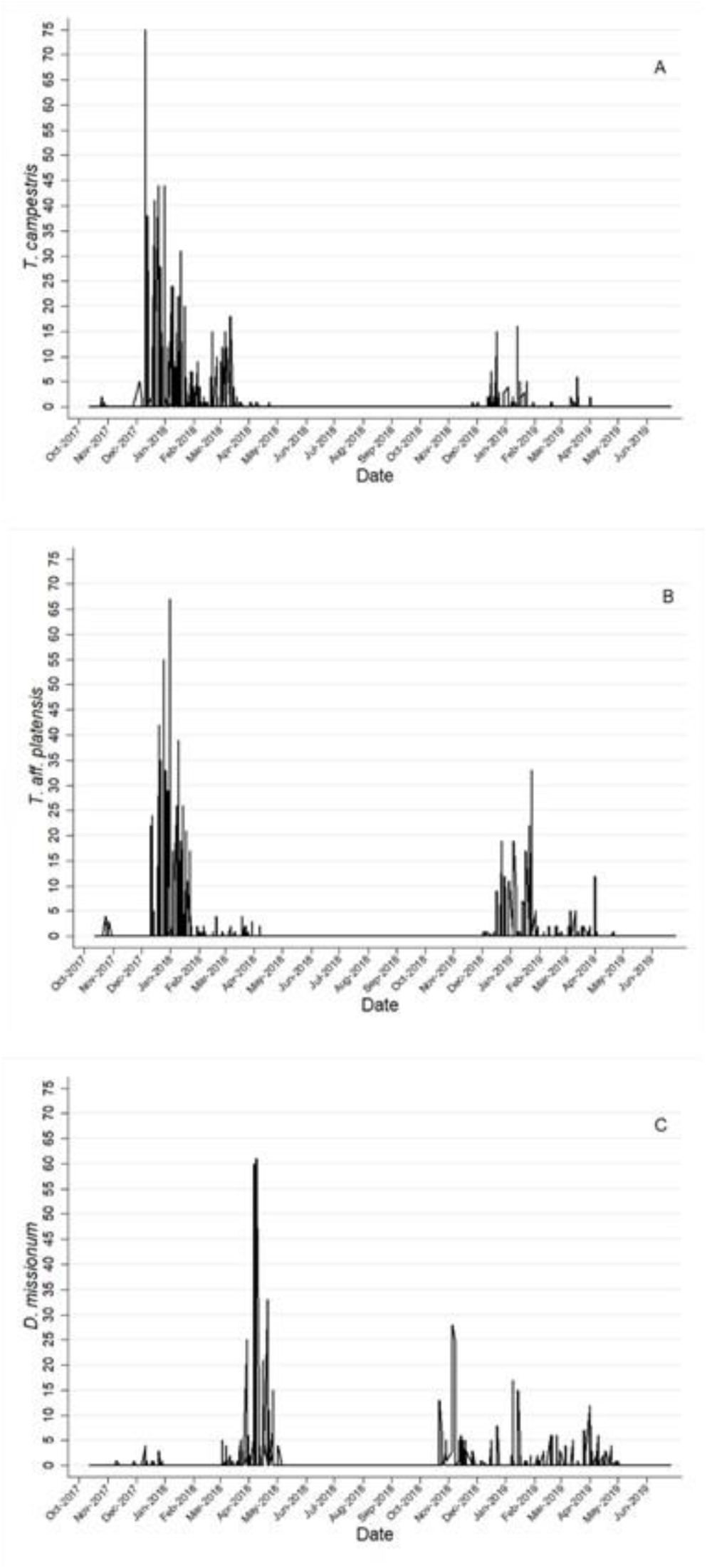
Seasonality of the 3 species most captured in the systematic collection: *T. campestris* (Fig. 2A), *T. aff. platensis* (Fig. 2B) and *D. missionum* (Fig. 2C).

The Nzi traps were responsible for 97.1% of the collections, while the Malaise traps captured 2.9%. In the “lowlands”, the Malaise trap captured 1.4% of the total individuals, while in the “highlands”, it was responsible for 21.7% of the collections. Of the 15 species captured in the systematic collections, the Nzi traps only did not capture *Tabanus pungens* Wiedemann, while the Malaise traps did not capture four of the 15 species: *P. quadripunctatus, Tabanus fuscofasciatus* Macquart, *Tabanus claripennis* (Bigot) and *Dasybasis ornatissima* (Brèthes).

Throughout the study period, the monthly average temperature ranged from 10.4 °C to 25.0 °C, the average minimum temperatures were between 6.0 °C and 19.4 °C, and the average maximum temperature ranged between 15.2 °C and 31.4 °C. The accumulated rainfall during this period was 2715.3 millimeters (mm), and the average relative humidity varied between 56% and 90% (Figure 3; Supplementary Table 1). During the first season (September 2017-May 2018), the accumulated rainfall was 944.8 mm, while in the second season (September 2018-May 2019), the accumulated rainfall was 1309.6 mm, marking a clear difference in rainfall, especially during the summer capture peak (December-January).

**Fig. 3.**
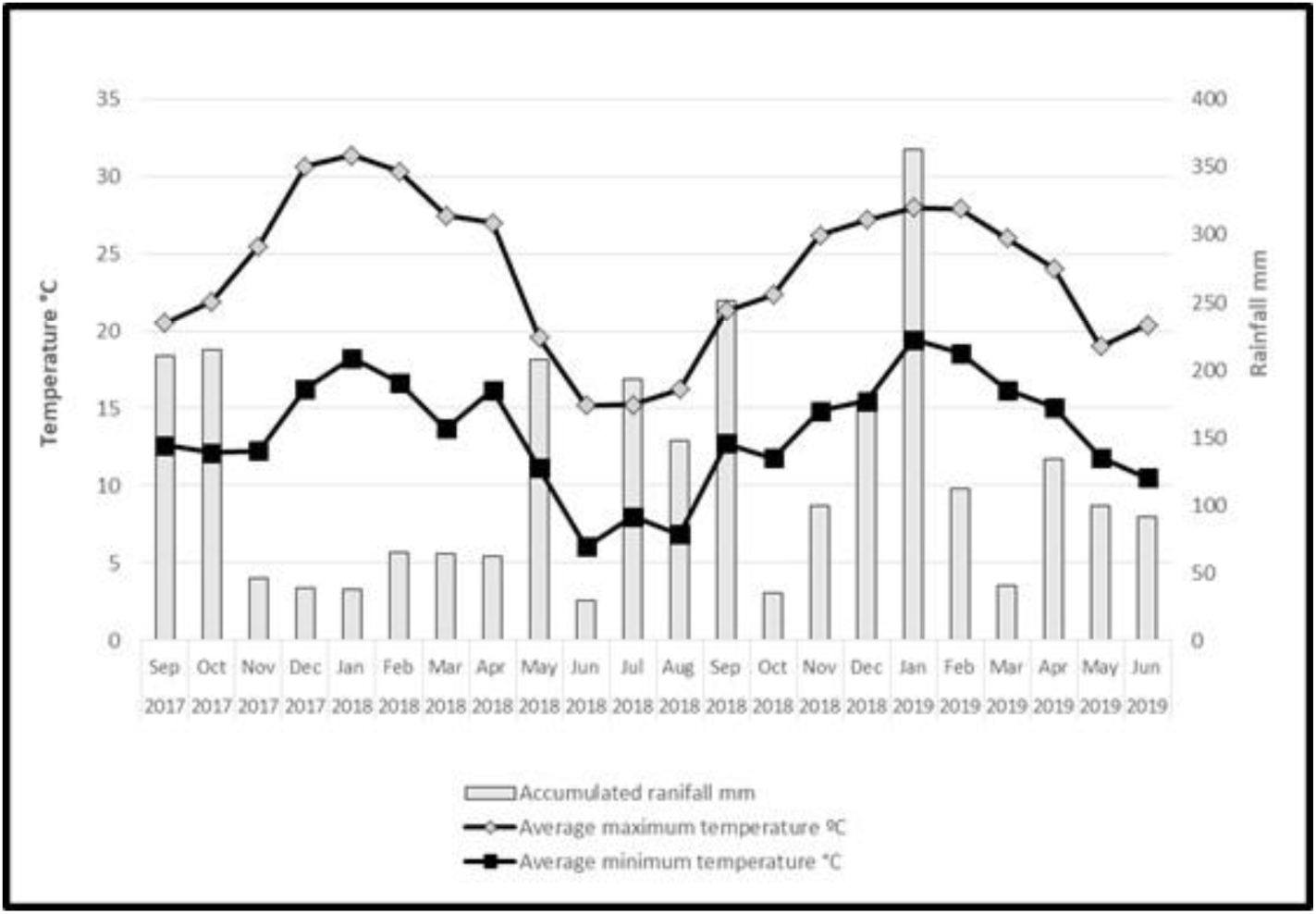
Monthly values of average maximum temperature, average minimum temperature and accumulated rainfall throughout the study period

Considering the response variable of the abundance of horse flies captured by the traps, during the entire study period, the average minimum and maximum temperatures, relative humidity and rainfall did not have a significant effect (p > 0.05) and were removed from the final model. The significant differences found were due to the increase in the mt and the environment. The mt (Chi_1;129_= 3,989.3, p<0.001) and environment (Chi_1;128_= 2,675.1, P<0.001) influenced the variation in the abundance of horseflies (Figure 4), and there was no interaction between these variables. According to the abundance models, there were 12 times more horseflies in the lowland environment than in the highland, and individuals were predicted to be present at average temperatures of 18 °C in the highland and 12 °C in the lowland. The models predicted approximately 200 individuals per trap at 25 °C in the lowland, while in the highland habitat, approximately 15 individuals were predicted.

**Fig. 4.**
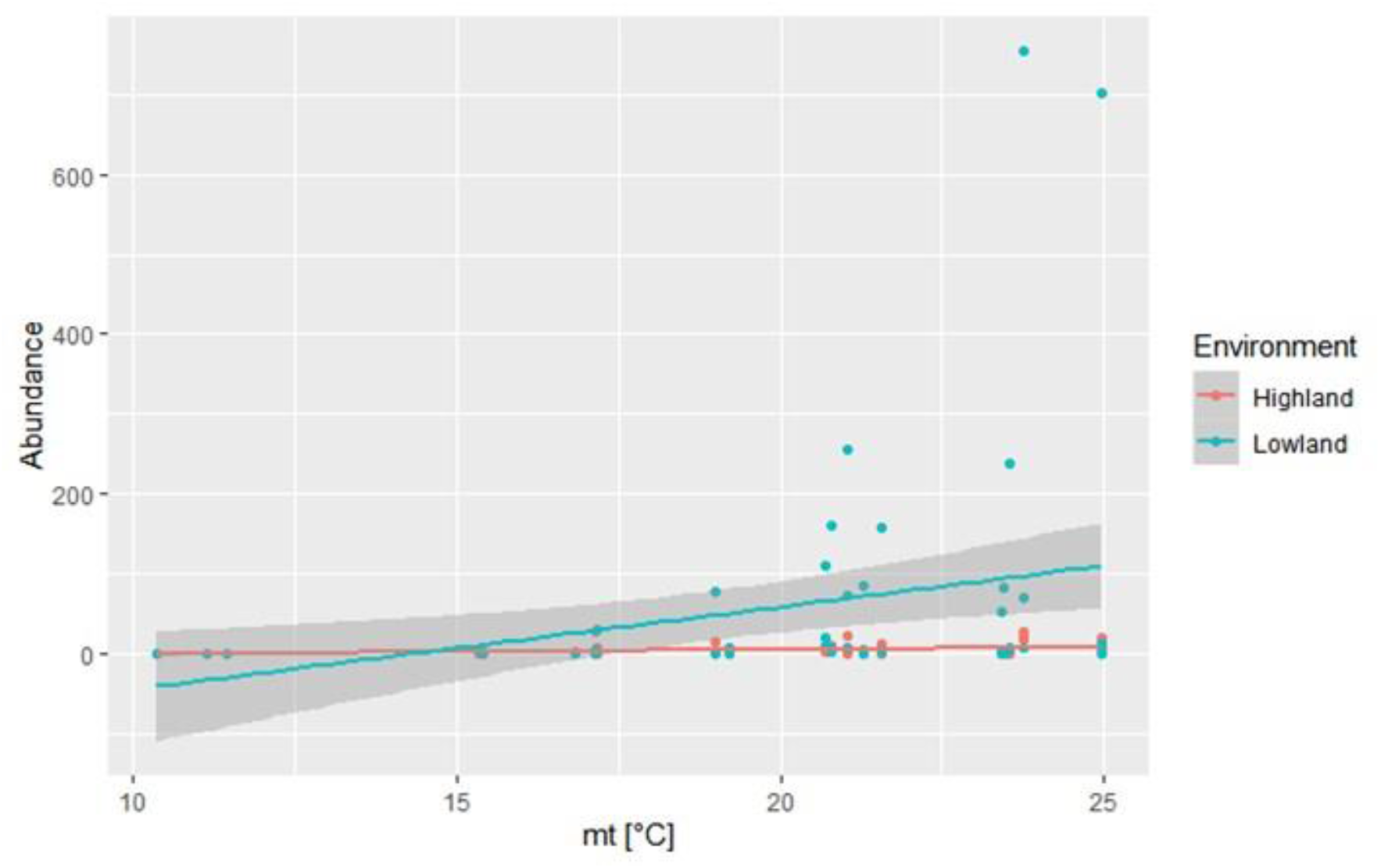
Abundance of horseflies in Tacuarembo, Uruguay, as a function of mean temperature (°C) (mt), month, and year according to GLM analysis with a quasi-Poisson distribution. The models for each environment are Abundance _Highland_ =exp ^− 6.886 +0.386*mt^ and Abundance _Lowland_ ^− 4.391+0.386*mt^

### Nonsystematic collections

A total of 455 individuals were collected in the departments of Colonia (n= 343), Tacuarembó (n= 74) and Paysandú (n= 38) (Figure 5; Table 3).

**Table 3.**
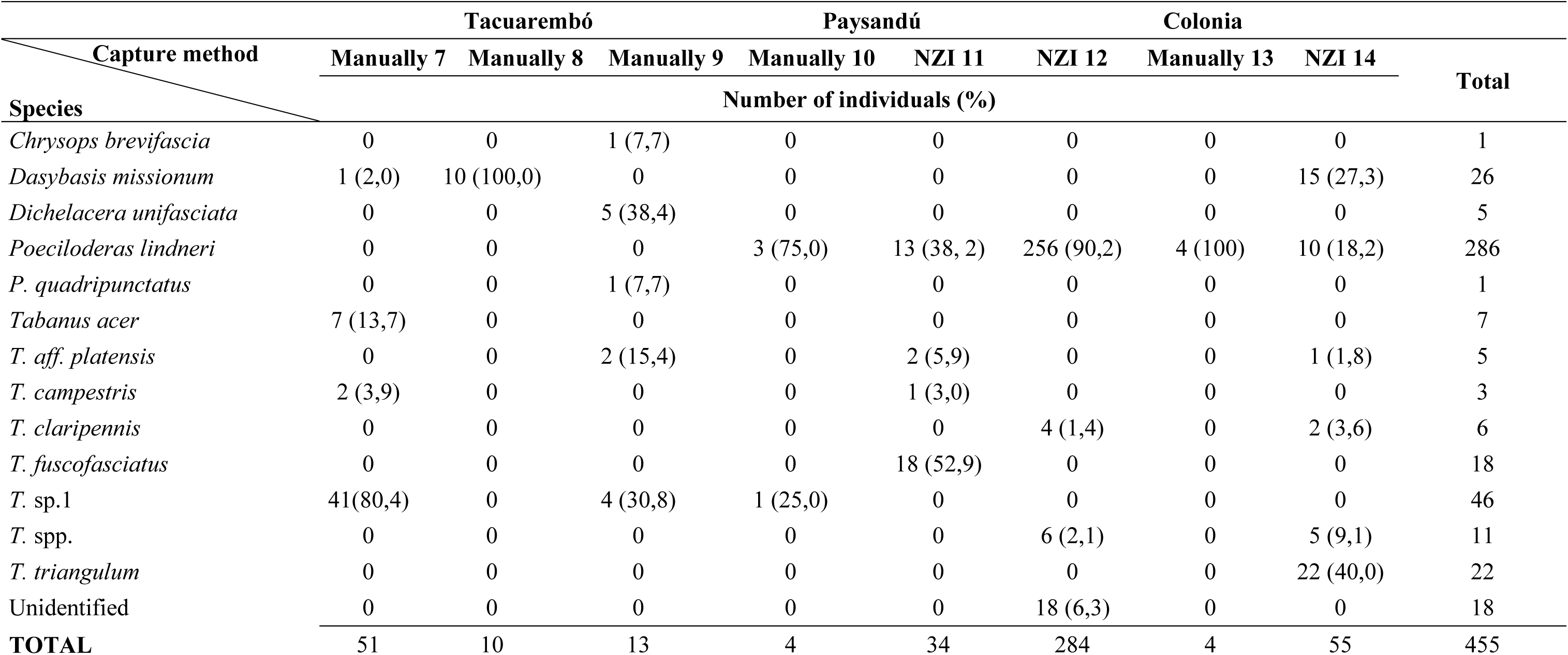
Number of individuals and species collected in nonsystematic collections using traps or manually in the departments of Paysandú, Colonia and Tacuarembó

**Fig. 5.**
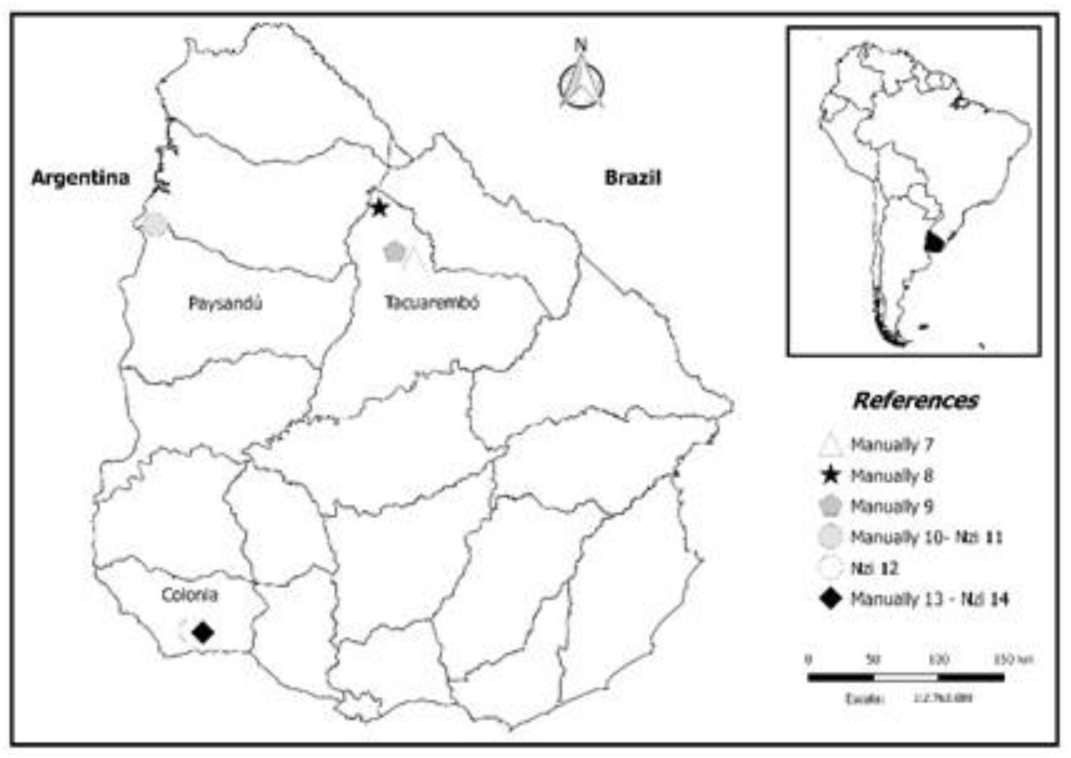
Location and capture method of nonsystematic collections.

In Colonia, 78.7% were *Poeciloderas lindneri* (Kröber), while in Paysandú, the two most captured species were *T. fuscofasciatus* (47.4%) and *P. lindneri* (42.1%). Regarding the manual captures in the Tacuarembó department, the most abundant species was the undescribed *Tabanus* sp.1 (60.8% of the captures).

Through this sampling, a total of 12 species were collected, with *Dichelacera unifasciata* Macquart being the only species not caught during the systematic captures. Eleven individuals classified as *Tabanus* spp. were also captured, but taxonomic identification was not achieved.

## Discussion

Of the 14 species taxonomically identified, *D. ornatissima* and *D. missionum* were recorded for the first time in Uruguay, although both species have also been recorded in Argentina, and *D. missionum* has been recorded in Brazil (Coscarón and Papavero 2009; Kruger and Krolow 2015). It is necessary to achieve the final identification of *T. aff. platensis* by comparison with “type” specimens. *Tabanus platensis* has not been reported in Uruguay, but its presence is expected, since its distribution in Argentina includes Chaco, Santa Fe, Entre Ríos and Buenos Aires (Coscarón and Papavero 2009).

Two of the three new records for Uruguay (*D. missionum* and *T. aff. platensis*) were among the three most prevalent species found during the systematic captures and had not been described as abundant in similar studies in southern Brazil (Kruger and Krolow 2015). An undescribed species (*Tabanus* sp.1) was also identified, which will be described posteriorly.

The fact that captures were concentrated between September and May, without horse fly activity during winter, differs from what was observed in southern Brazil, where, despite a marked decrease in captures during winter, the presence of horse flies still occurred (Kruger and Krolow 2015). In southern Brazil, there is a peak in horse fly abundance during spring and a gradual decrease as winter approaches, but it is still possible to capture horse flies during this season (Kruger and Krolow 2015). In the Tacuarembó department, which belongs to the same Pampa biome as southern Brazil, the largest number of horse flies was observed during summer, with lower peaks during spring and autumn. This variation can be explained by climatological differences between zones or between years as well as differences in the species of horse flies captured in the two regions (Burnett and Hays 1974; Gorayeb 1995; Kruger and Krolow 2015). In Tacuarembó, the peak recorded in autumn was mostly due to *D. missionum*. This peak was not recorded in Rio Grande do Sul, where *D. missionum* was not among the most abundant species (Kruger and Krolow 2015). It is important to note that different species respond differently to the variation in climatic factors (Van Hennekeler et al. 2008).

Throughout the study period, only the mean temperature and environment had statistically significant effects on the number of individuals captured. In contrast, some studies have related the abundance of horse flies with an increase in relative humidity (Cardenas et al. 2012) interacting with the mean temperature (Kruger and Krolow 2015) and highlight that the influence of this variable differs according to species (Van Hennekeler et al. 2008; Kruger and Krolow 2015).

The environment was a determining factor related to the number of horse flies collected. The traps located in “lowland” habitats, with native forest and creeks, captured 12 times more horse flies on average than the traps located in the “highland” habitat. These findings would allow a more efficient use of traps as a measure to mitigate horse fly attack on commercial farms through their use in environments with a greater number of horse flies. In addition, grazing management patterns that preclude the animals from grazing in the most infested paddocks during population peaks could be established to prevent the hosts and parasites from being simultaneously present (Foil and Hogsette 1994; Baldacchino et al. 2014). Barros (2001) did not report relevant differences in the number of horse flies in the Pantanal biome (Brazil) when comparing captures carried out in areas of native forests with captures carried out in grasslands. However, Gorayeb (2000) found differences in the abundance and diversity of species in different environments in Santa Barbara, Pará (Brazil), after performing captures in two environments: near forest and in areas of clear fields (grassland). This author captured 71.5% of the specimens in the environment near the forest and 28.5% in the clear field environment. In addition, 23 of the 47 species were present in both environments, while 14 were exclusively found in the environment near forest, and 9 were only found in the grassland environment.

Even when the first season of captures occurred during a dry summer and the second during a rainy summer, a similar pattern in population peaks was observed. However, during the second season of systematic captures, a remarkable decrease was observed in the number of individuals caught. This decrease in captures during the second season may have been influenced by the constant presence of the traps; however, no remarkable decrease in the density of horse fly populations in the environment has been evident in previous studies using traps as a control method (Vale et al. 1988; Foil and Hogsette 1994; Baldacchino et al. 2014). Another possibility is that this decrease was related to climatological and/or environmental factors, such as the amount of rainfall during the second year, which could affect the abundance and activity of the horse flies, as well as trap efficiency.

The variations in the prevalence and the species captured in different regions denote the need to continue the collection of horse flies in different areas of the country. As an example, in the department of Colonia, the most prevalent species found was *P. lindneri*, while in the systematic collections performed in the department of Tacuarembó, this species accounted for less than one percent of the total captures. Additionally, Coscarón and Martínez (2019) highlighted the importance of carrying out new captures in Uruguay, especially in rural areas, to assess whether the diversity of species reported is still maintained after urbanization.

Horse flies are detrimental to livestock production due to their direct effects on animals and their ability to act as vectors for at least 35 pathogens (Krinsky 1976, Foil 1989, Baldacchino et al. 2014). Relevant agents of disease potentially transmissible by horse flies in Uruguay include *A. marginale*, bovine leucosis virus, *Brucella abortus, Leptospira* spp, and equine infectious anemia virus. More studies need to be performed to determine the role of the Tabanidae species found in Uruguay on the epidemiology of these agents.

## Acknowledgments

We would like to acknowledge Marcelo Alfonso for help with the artwork preparation and Gonzalo Escayola for helping with the data collection.

---

**Supplementary table 1.**
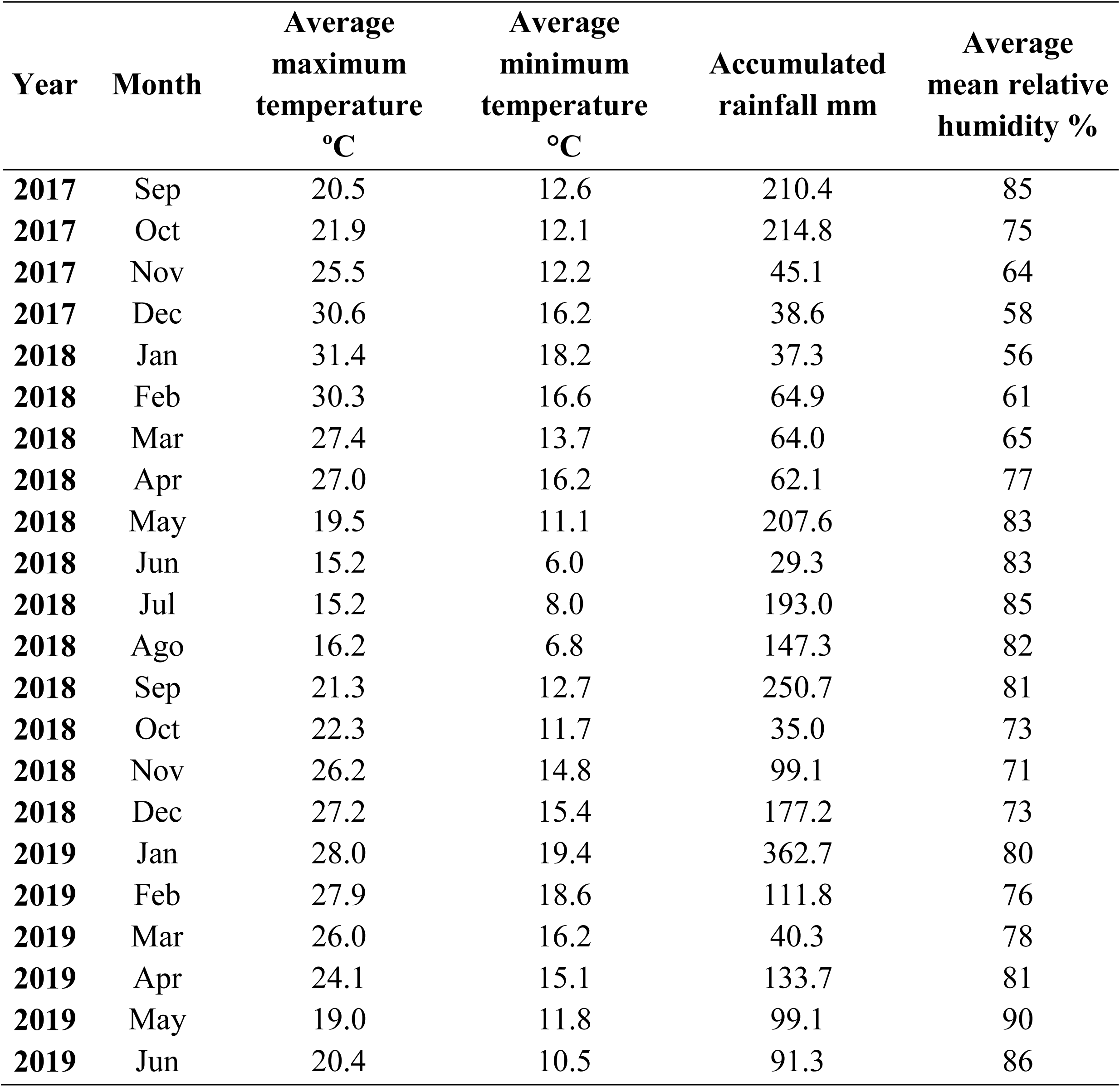
Climatic variables that were recorded and evaluated throughout the study period.

| **Year** | **Month** | **Average**<br>**maximum**<br>**temperature**<br>**°C** | **Average**<br>**minimum**<br>**temperature**<br>**°C** | **Accumulated**<br>**rainfall mm** | **Average**<br>**mean relative**<br>**humidity %** |
| --- | --- | --- | --- | --- | --- |
| **2017** | Sep | 20.5 | 12.6 | 210.4 | 85 |
| **2017** | Oct | 21.9 | 12.1 | 214.8 | 75 |
| **2017** | Nov | 25.5 | 12.2 | 45.1 | 64 |
| **2017** | Dec | 30.6 | 16.2 | 38.6 | 58 |
| **2018** | Jan | 31.4 | 18.2 | 37.3 | 56 |
| **2018** | Feb | 30.3 | 16.6 | 64.9 | 61 |
| **2018** | Mar | 27.4 | 13.7 | 64.0 | 65 |
| **2018** | Apr | 27.0 | 16.2 | 62.1 | 77 |
| **2018** | May | 19.5 | 11.1 | 207.6 | 83 |
| **2018** | Jun | 15.2 | 6.0 | 29.3 | 83 |
| **2018** | Jul | 15.2 | 8.0 | 193.0 | 85 |
| **2018** | Ago | 16.2 | 6.8 | 147.3 | 82 |
| **2018** | Sep | 21.3 | 12.7 | 250.7 | 81 |
| **2018** | Oct | 22.3 | 11.7 | 35.0 | 73 |
| **2018** | Nov | 26.2 | 14.8 | 99.1 | 71 |
| **2018** | Dec | 27.2 | 15.4 | 177.2 | 73 |
| **2019** | Jan | 28.0 | 19.4 | 362.7 | 80 |
| **2019** | Feb | 27.9 | 18.6 | 111.8 | 76 |
| **2019** | Mar | 26.0 | 16.2 | 40.3 | 78 |
| **2019** | Apr | 24.1 | 15.1 | 133.7 | 81 |
| **2019** | May | 19.0 | 11.8 | 99.1 | 90 |
| **2019** | Jun | 20.4 | 10.5 | 91.3 | 86 |

## References

Baldacchino F, Desquesnes M, Mihok S, Foil L, Duvallet G, Jittapalapong S (2014) Tabanids: Neglected subjects of research, but vectors of disease agents Important!. Infect Genet Evol 28: 596–615. http://dx.doi.org/10.1016/j.meegid.2014.03.029

Barros ATM (2001) Seasonality and relative abundance of Tabanidae (Diptera) captured on horses in the Pantanal, Brazil. Mem Inst Oswaldo Cruz 96: 917–923. http://dx.doi.org/10.1590/S0074-02762001000700006

Burnett AM, Hays KL (1974) Some influences of meteorological factors on flight activity of female horse flies (Diptera: Tabanidae). Environ Éntomol 3: 515–521. https://doi.org/10.1093/ee/3.3.515

Cardenas RE, Hernandez NL, Barragán AR, Dangles O (2012) Differences in morphometry and activity Among tabanid fly assemblages in an Andean tropical montane cloud forest: indication of altitudinal migration?. Biotropica 45: 63–72. https://doi.org/10.1111/j.1744-7429.2012.00885.x

Catálogo Taxonômico da Fauna do Brasil (2019) http://fauna.jbrj.gov.br/fauna/listaBrasil/ConsultaPublicaUC/ConsultaPublicaUC. do Accessed 04 August 2019.

Chvála M, Lyneborg L, Moucha J (1972) The Horse Flies of Europe (Diptera, Tabanidae). Entomological Society of Copenhagen, Copenhagen. ISBN: 978-09-00-84857-5

Coscarón S, Martinez M (2019) Checklist of tabanidae (Insecta: Diptera) from Uruguay. J Éntomol Soc Argentina 78: 40–46. https://doi.org/10.25085/rsea.780105

Coscarón S, Papavero N (2009) Catalogue of Neotropical Diptera. Tabanidae. Neotropical Diptera 16: 1–199.

Coscarón S (1998) Tabanidae (Diptera-Insecta). In: Morrone JJ, Coscarón S (ed) Biodiversidad de artrópodos argentinos: una perspectiva biotaxonómica Sur, La Plata, pp 341–352

Foil LD (1989) Tabanids as vectors of disease agents. Parasitol Today 5: 88–96. https://doi.org/10.1016/0169-4758(89)90009-4

Foil LD, Hogsette JA (1994) Biology and Control of Tabanids, stable flies and horn flies. Rev Sci Tech 13: 1125–1158.

Gorayeb IS (1995) Tabanidae (Diptera) da Amazônia. XI. Sazonalidade espécies das da Amazônia Oriental e correlação com fatores climáticos. Bol Mus Paraense Emílio Goeldi 9: 241–281. http://repositorio.museu-goeldi.br:8080/jspui/handle/mgoeldi/1004

Gorayeb IS (2000) Tabanidae (Diptera) da Amazônia. XVI – Atividade diurna de hematofagia de espécies da Amazônia Oriental, em áreas de mata e pastagem, correlacionada com fatores climáticos. Bol Mus Paraense Emílio Goeldi 16: 23–63. http://repositorio.museu-goeldi.br:8080/jspui/handle/mgoeldi/1021

Hawkins JA, Love JN, Hidalgo RJ (1982) Mechanical transmission of Anaplasmosis by tabanids (Diptera, Tabanidae). Am J Vet Res 43: 732–734.

Henriques AL, Krolow TK, Rafael JA (2012) Corrections and additions to Catalog of Neotropical Diptera (Tabanidae) of Coscarón & Papavero (2009). Revista Brasileira de Entomologia 56: 277–280. http://dx.doi.org/10.1590/S0085-56262012005000042

Hornok S, Földvári G, Elek V, Naranjo V, Farkas R, de la Fuente J (2008) Molecular identification of *Anaplasma marginale* and rickettsial endosymbionts in bloodsucking flies (Diptera: Tabanidae, Muscidae) and hard ticks (Acari: Ixodidae). Vet Parasitol 54: 354–359. https://doi.org/10.1016/j.vetpar.2008.03.019

INIA (2019). GRAS INIA data. http://www.inia.uy/gras/Clima/Banco-datos-agroclimatico).

Krinsky WL (1976) Animal-disease agents transmitted by horse flies and deer flies (Diptera, Tabanidae). J Med Éntomol 13: 225–275. https://doi.org/10.1093/jmedent/13.3.225

Krüger RF, Krolow TK (2015) Seasonal patterns of horse fly richness and abundance in the Pampa biome of southern Brazil. J Ecol Vector 40: 364–372. https://doi.org/10.1111/jvec.12175

Magnarelli LA, Anderson JF (1980) Feeding-behavior of Tabanidae (Diptera) on cattle and serological analysis of partial blood meals. Environ Éntomol 9: 664–667. https://doi.org/10.1093/ee/9.5.664

Mihok S (2002) The development of a multipurpose trap (the Nzi) for tsetse and other biting flies. Bull Éntomol Res 92: 385–403. https://doi.org/10.1079/BER2002186

Miraballes C, Lucas M, Krolow T, Riet-Correa F, Medeiros de Barros AT, Kruger R, Saravia A (2019) data for Diversity and seasonality of horse flies (Diptera: Tabanidae) in Uruguay”, Mendeley Data, v2 http://dx.doi.org/10.17632/zstbdddgv8.2

Perich MJ, Wright RE, Lusby KS (1986) Impact of horse flies (Diptera: Tabanidae) on beef-cattle. J Econ Éntomol 79: 128–131. https://doi.org/10.1093/jee/79.1.128

Scoles G, Miller A, Foil L (2008) Comparison of the Efficiency of Biological Transmission of *Anaplasma marginale* (Rickettsial: Anaplasmataceae) by *Dermacentor andersoni* Stiles (Acari: Ixodidae) With Mechanical Transmission by the Horse Fly, Tabanus fuscicostatus Hine (Diptera: Muscidae). J Méd Éntomol 45: 109–114. https://doi.org/10.1093/jmedent/45.1.109

StataCorp. (2015) Stata Statistical Software: Release 14. College Station, TX: StataCorp LP.

Thompson, PH (1969) Methods for Collecting Tabanidae (Diptera) Ann Éntomol Soc Am 62: 50–57. https://doi.org/10.1093/aesa/62.1.50

Thorsteinson AJ, Bracken GK, Hanec W (1965) The orientation of horse flies and deer flies (Tabanidae, Diptera). III. The use of traps in the study of orientation of Tabanids in the field. Éntomol Exp Appl 8: 189–192. https://doi.org/10.1111/j.1570-7458.1965.tb00853.x

Vale GA, Lovemore DF, Flint S, Cockbill GG (1988) Odour-baited targets to control tsetse flies, *Glossina* spp. (Diptera: Glossinidae), in Zimbabwe. Bull Éntomol Res 78: 31–49. https://doi.org/10.1017/S0007485300016059

Van Hennekeler K, Jones RE, Skerratt LF, Fitzpatrick LA, Reid SA, Bellis GA (2008). A comparison of trapping methods for Tabanidae (Diptera) in North Queensland, Australia. Méd Vet Éntomol 22: 26–31. https://doi.org/10.1111/j.1365-2915.2007.00707.x

